# β-cyclocitral is a conserved root growth regulator

**DOI:** 10.1101/337162

**Authors:** Alexandra J. Dickinson, Kevin Lehner, Jianing Mi, Kun-Peng Jia, Medhavinee Mijar, José Dinneny, Salim Al-Babili, Philip N. Benfey

**Affiliations:** Department of Biology, Duke University, Durham, NC 27708; Howard Hughes Medical Institute, Duke University, Durham, NC 27708; King Abdullah University of Science and Technology (KAUST), Biological and Environmental Sciences and Engineering Division, The Bioactives Lab, Thuwal, 23955-6900, Saudi Arabia.; Department of Plant Biology, Carnegie Institute of Science, Stanford, CA 94305; Department of Biology, Stanford University, Palo Alto, CA 94305

## Abstract

Natural compounds capable of increasing root depth and branching are desirable tools for enhancing stress tolerance in crops. We devised a sensitized screen to identify natural metabolites capable of regulating root traits in Arabidopsis. β-cyclocitral, an endogenous root compound, was found to promote cell divisions in root meristems and stimulate lateral root branching. β-cyclocitral rescued meristematic cell divisions in *ccd1ccd4* biosynthesis mutants and β-cyclocitral-driven root growth was found to be independent of auxin, brassinosteroid, and ROS signaling pathways. β-cyclocitral had a conserved effect on root growth in tomato and rice and generated significantly more compact crown root systems in rice. Moreover, β-cyclocitral treatment enhanced plant vigor in rice plants exposed to salt-contaminated soil. These results indicate that β-cyclocitral is a broadly effective root growth promoter in both monocots and eudicots and could be a valuable tool to enhance crop vigor under environmental stress.

**One Sentence Summary:** β-cyclocitral is a metabolite of β-carotene that was identified using a sensitized chemical screen and acts broadly across plants to enhance root growth and branching.

## Introduction

A rapidly increasing world population, coinciding with changes in climate, creates a need for new methods to stabilize and improve crop productivity under harsh environmental conditions. Exogenously applied metabolites and phytohormones, such as auxin, cytokinin, and ethylene, have had profound impacts on agriculture by selectively killing weeds, promoting shoot growth, and optimizing fruit ripening (1–5). Root traits such as growth and branching are appealing targets for enhancing plant performance, due to their essential role in nutrient and water uptake. In Arabidopsis, root development begins with the formation of a primary root during embryogenesis. Root branching is initiated by oscillations in gene expression at the tip of the primary root (6). These oscillations establish the future position of *de novo* roots called lateral roots (LRs) and are referred to as the “LR clock.” Stereotyped cell divisions promote LR primordia development by increasing cell number and generating all root cell types (7). In both primary and LRs, growth is maintained through cell divisions in the meristem and the subsequent elongation of daughter cells. Previously, inhibition of the carotenoid pathway was shown to reduce root growth and branching in Arabidopsis, suggesting that this pathway is important for LR development (8).

The carotenoid pathway is a rich source of metabolites, called apocarotenoids, several of which are known regulators of root development (Figure S1) (9, 10). Strigolactones and abscisic acid (ABA), which are apocarotenoid phytohormones, regulate root growth and LR branching, respectively (11–13). Previously, an inhibitor of carotenoid cleavage dioxygenases (CCDs) called D15 was found to decrease LR branching through an ABA and strigolactone independent-mechanism in Arabidopsis (8). This suggested one or more unidentified apocarotenoids could be positive regulators of LR development. Isolating these compounds through genetic means is difficult because each CCD has multiple substrates and produces a variety of different compounds, making it impossible to selectively eliminate a single apocarotenoid. Therefore, to identify new natural compounds capable of promoting root growth and development, we leveraged D15 to characterize the effects of exogenously applied apocarotenoids on root traits in a sensitized background.

## Results

To identify apocarotenoids involved in LR development we utilized a targeted chemical genetic approach to screen apocarotenoids for their ability to enhance root branching in the presence of D15. The maximum concentration of D15 tested (100 μM) decreased primary root length and completely inhibited LR capacity, the number of LRs that emerge after excision of the primary root apical meristem (8). By titrating D15 to 30 μM, the concentration at which it decreases LR emergence by 50% (IC_50_), we could measure changes in LR capacity with enhanced sensitivity (Figure 1A). Most apocarotenoids tested, including ABA and GR24, a synthetic strigolactone analogue, further decreased LR capacity when combined with D15 (Figure S2). Two apocarotenoids, dihydroactinidiolide (DHAD) and β-cyclocitral, were found to increase LR branching in the presence of D15. These compounds were previously found to trigger the reactive oxygen species response and increase leaf tolerance to high light stress (14–16).

**Figure 1.**
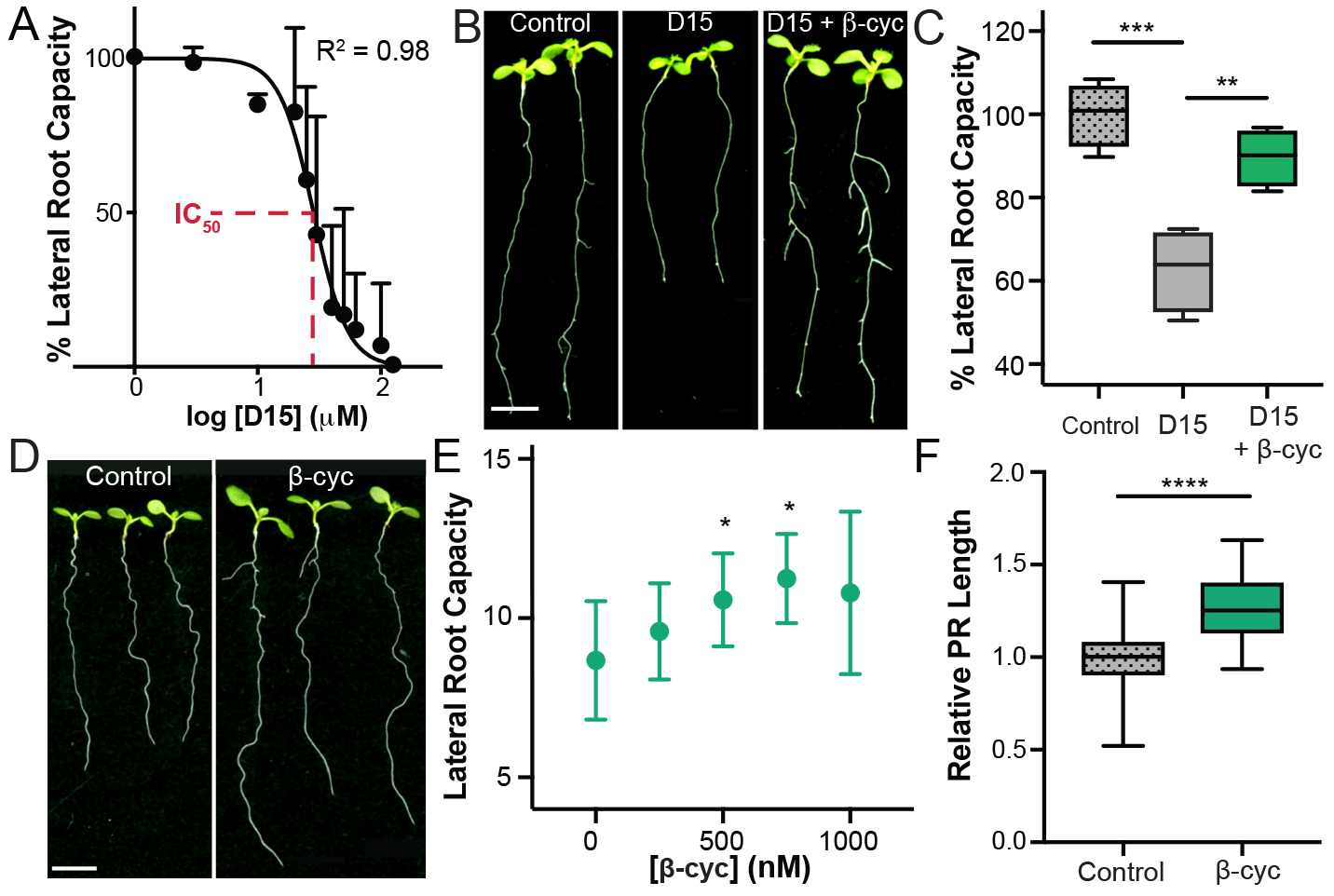
Identification of β-cyclocitral, a root growth promoter. A) The LR capacity of D15-treated plants, normalized to control plants. The IC_50_ is highlighted in red. B) Seedlings after treatment with D15 and β-cyclocitral (β-cyc) (scale bar = 5 mm) C) LR capacity of plants treated with D15 and β-cyclocitral. D) Arabidopsis seedlings treated with 750 nM β-cyclocitral (scale bar = 5 mm). E) Quantification of LR capacity of seedlings treated with increasing concentrations of β-cyclocitral. F) Quantification of primary root length in β-cyclocitral-treated plants. The symbols *, **, ***, and **** indicate p values ≤ 0.05, 0.01, 0.001, and 0.0001, respectively.

To further explore the effects of DHAD and β-cyclocitral on LR branching, we varied their concentrations in the presence of D15 at its IC_50_ concentration. We found that even the most effective concentration of DHAD only increased root branching by 14%. However, application of volatile β-cyclocitral had a much more dramatic effect, promoting root branching by nearly 40% (Figure 1B-C). Therefore, we focused on β-cyclocitral’s mechanism of action. β-cyclocitral has been identified endogenously in dozens of plant species, including tomato (17), rice (18), parsley (19), tea (*20*), grape (*21*), various trees (22, 23), and moss (24), indicating that its presence in plants is evolutionarily conserved. We further identified endogenous β-cyclocitral in Arabidopsis and rice roots at levels of 0.097 and 0.47 ng/mg dry weight, respectively, using HPLC-MS (Figure S3). A gene ontology (GO) term enrichment analysis of previously published data in Arabidopsis revealed that genes upregulated by β-cyclocitral are important for the immune system, metabolite catabolism, and abiotic stress responses, indicating that β-cyclocitral may play a number of roles in growth and development (Table S1) (13).

To determine the effects of exogenous β-cyclocitral application in the absence of D15, we examined changes in root architecture upon treatment with β-cyclocitral alone. This treatment enhanced primary root length and LR branching by 30% as compared to control treatment (Figure 1D-F). The maximum efficacy of exogenous β-cyclocitral treatment occurred at a concentration of 750 nM, which is comparable to the levels at which exogenous ABA and strigolactones confer phenotypic changes (13). Apocarotenoids with nearly identical chemical structures, such as dimethyl-β-cyclocitral and β-ionone, did not increase root branching (Figure S1-2). These results suggest that β-cyclocitral is the active molecule or is a unique precursor to the active metabolite regulating LR branching (25).

To understand how β-cyclocitral promotes LR branching, we characterized its effect on LR development in the presence and absence of D15. Its ability to increase LR capacity suggests that it either increases initiation of new LRs or induces LR outgrowth. Since initiation is preceded by the LR clock, we monitored*pDR5:LUC*, a marker line that gives a readout of LR clock oscillations, after D15 and β-cyclocitral treatment. The region of the root tip that experiences the peak luminescence oscillation intensity becomes competent to form LR primordia (Figure S4) (6). In D15-treated roots, the peak oscillation intensity was significantly lower as compared to that of control roots (Figure 1D-E). β-cyclocitral did not affect the peak oscillation intensity, with or without D15, indicating that it does not increase initiation of LR primordia (Figure S4). To further test this, we examined the effect of β-cyclocitral on the formation of the first cell divisions in LR primordia. As expected, the IC_50_ concentration of D15 decreased the number of initiated LR primordia by approximately 50% compared to untreated plants, as reported by the*pWOX5:GFP* marker line (Figure S5A)(26). Consistent with its inability to restore LR clock amplitude, β-cyclocitral did not increase the number of WOX5^+^ primordia in D15-treated plants. Additionally, it did not increase WOX5^+^ sites in the absence of D15. These results suggest that β-cyclocitral does not have a role in determining the number of initiated LR primordia, and must instead promote LR branching by stimulating developmental stages that occur after initiation. To test this hypothesis, we used an EN7 marker line (*pEN7:GAL4; UAS:H2A-GFP*), which reports formation of the endodermis, an intermediate step before primordia emergence (27). β-cyclocitral doubled the number of EN7+ primordia in D15-treated plants and increased EN7^+^ primordia by 16% as compared to untreated plants (Figure S5B). Taken together, our results indicate that β-cyclocitral does not affect the total number of initiated LR primordia, but instead enhances subsequent pre-emergence growth of LR primordia.

To further determine how β-cyclocitral stimulates root growth and branching, we quantified its effect on the developmental stages in primary roots. Because β-cyclocitral stimulates progenitor cell divisions in LR primordia prior to cell elongation, we hypothesized that it induces root growth by increasing cell divisions in the root meristem. To test this, we examined meristematic cell numbers and cell elongation in primary roots treated with β-cyclocitral. The number of meristematic cells increased more than 20% upon treatment with β-cyclocitral, while cell elongation remained unchanged (Figure 2A-B, Figure S6). This suggests that β-cyclocitral enhances divisions in undifferentiated cells in the root meristem. To further test this, we characterized levels of a cyclin-dependent kinase in the meristems of plants harboring the construct, *pCYCB1;1:CYCB1;1-GFP* (Figure 2C-D). We found that β-cyclocitral significantly increased the number of GFP positive cells in the root meristem. These results indicate that β-cyclocitral induces root growth by stimulating meristematic cell divisions.

**Figure 2.**
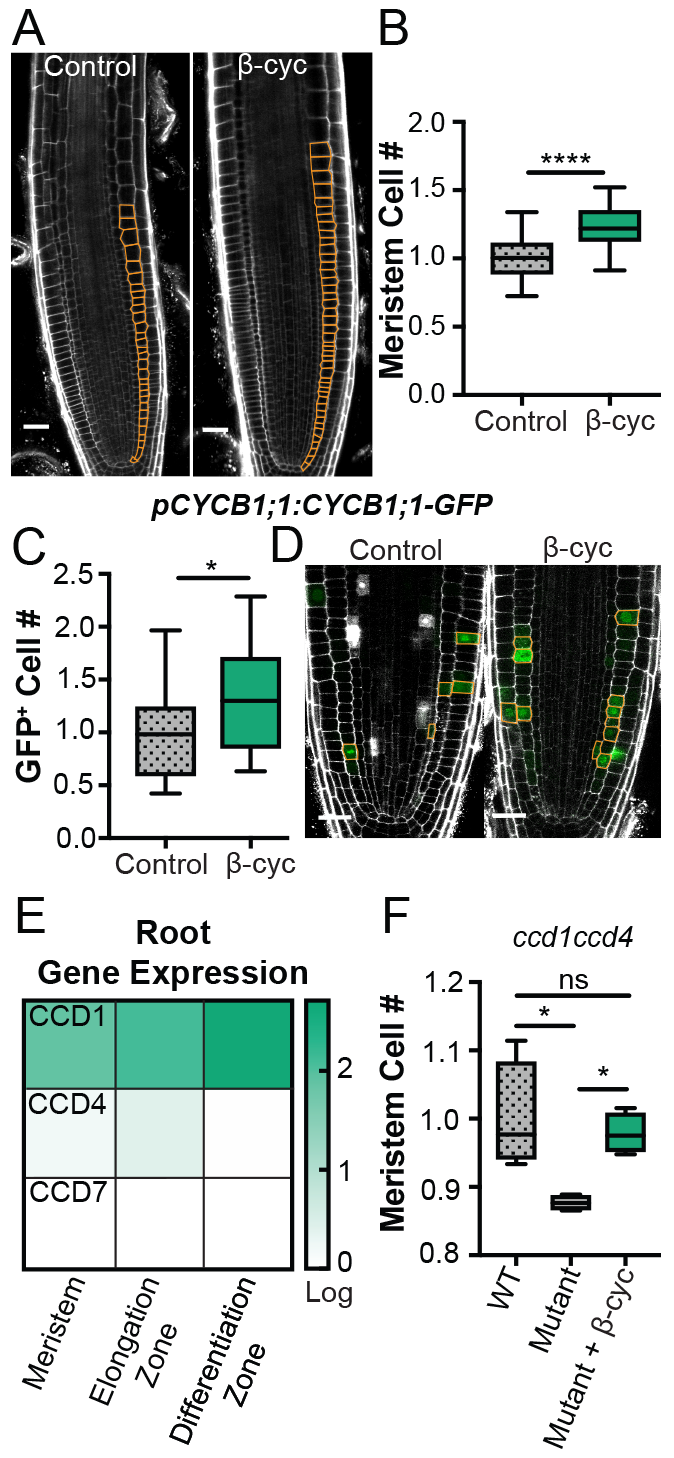
β-cyclocitral induces meristematic cell divisions. A) Confocal images of primary root meristems (scale bar = 50 μm). Meristematic cortex cells are highlighted in orange. B) Relative number of cortex cells in the primary root meristems of treated and control plants. C) Relative number of GFP positive cells in the root meristem of *pCYCB1;1:CYCB1;1-GFP* seedlings treated with β-cyclocitral. D) Confocal images of root meristems in *pCYCB1;1:CYCB1;1-GFP* seedlings (scale bar = 25 μm). GFP positive cells are outlined in orange. E) CCD1, CCD4, and CCD7 gene expression (log[3xFPKM]) in the three developmental zones at the root tiμ F) Relative number of cortex cells in the primary root meristems of WT and *ccd1ccd4* double mutants with and without β-cyclocitral. The symbols * and **** indicate p values ≤ 0.05 and 0.0001, respectively.

To further characterize endogenous β-cyclocitral, we investigated the role of CCD1, CCD4, and CCD7, which cleave β-carotene and may therefore contribute to the formation of β-cyclocitral. Previous work indicated that CCD1 and CCD4 are expressed in the root meristem and elongation zone, but CCD7 is not expressed in the meristem, elongation zone, or beginning of the differentiation zone at the root tip (Figure 2E) (28, 29). To characterize the effect of enzymatically depleting β-cyclocitral in roots, we generated a *ccd1ccd4* double mutant. This mutant had significantly fewer meristematic cells as compared to wild-type roots (Figure 2F). Meristem cell number could be rescued upon application of β-cyclocitral. These data further indicate that β-cyclocitral has an endogenous regulatory role in root development. To determine if β-cyclocitral acts through previously characterized auxin, ROS, or brassinosteriod pathways, which induce meristematic divisions, we quantified the effect of β-cyclocitral on hormone-responsive marker lines and mutants. The auxin-responsive lines *pDR5:GFP*, *pPIN3:PIN3-GFP*, *pPLT2:CFP*, and *pPLT2:PLT2-GFP* all showed significantly increased meristematic divisions upon β-cyclocitral treatment, yet none had changes in meristem fluorescence (Figure S7). Moreover, inhibition of auxin transport using NPA did not affect the ability of β-cyclocitral to increase root length (Figure S8). We also found no defects in β-cyclocitral induction of root growth in mutants in ROS (*upb1-1*) and brassinosteroid (*bri1-4*) signaling pathways (Figure S8), suggesting that β-cyclocitral does not regulate meristem divisions through the major pathways that have previously been shown to stimulate meristem growth.

To determine if β-cyclocitral has an effect on agriculturally important plant species, we assayed root growth in tomato and rice seedlings and found that β-cyclocitral significantly increased primary and LR length in *Solanum lycopersicum* (Figure 3A, Figure S9) and in rice seedlings. This provides strong evidence that β-cyclocitral is a conserved regulator able to stimulate primary and LR length and meristem size in eudicots and monocots (Figure 3B). In addition, β-cyclocitral reduced the angle between LRs and the primary root - generating steeper root systems - in all three plant species.

**Figure 3.**
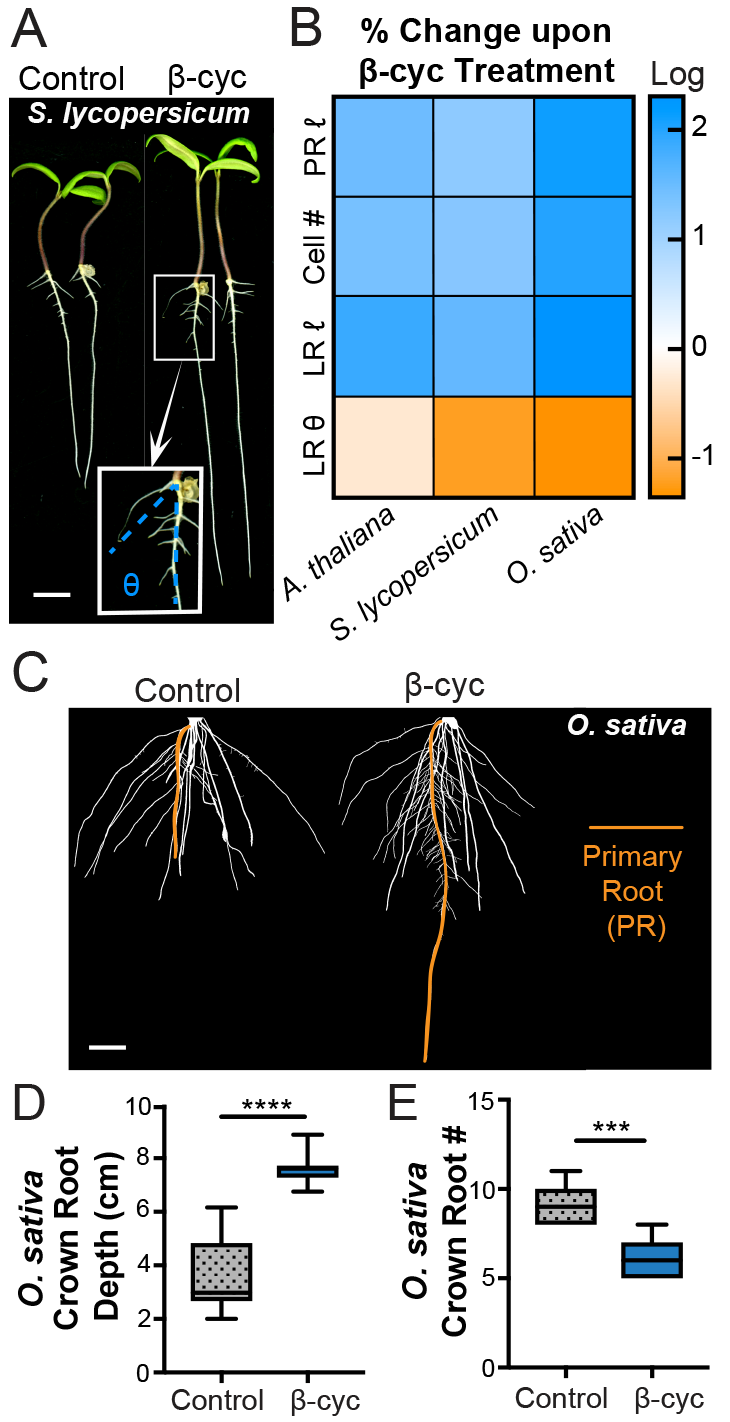
β-cyclocitral has conserved effects on root architecture. A) Tomato seedlings treated with β-cyclocitral (scale bar = 10 mm). Inset: the growth angle (θ) between the tip of the LR and the primary root measured to quantify the steepness of LRs is shown in blue. B) Heat map depicting the increase (blue) or decrease (orange) in primary root length (PR *ℓ*), meristematic cell number (Cell #), LR length (LRl *ℓ*), and angle of LR growth (LR θ) upon treatment with β-cyclocitral in Arabidopsis, tomato, and rice. C) Root systems of 9311 rice seedlings treated with β-cyclocitral (scale bar = 10 mm). The primary roots are highlighted in orange. D) Quantification of the average depth of the crown roots in rice. E) Quantification of the number of crown roots per seedling. The symbols *** and **** indicate p values ≤ 0.001 and 0.0001, respectively.

β-cyclocitral had a striking ability to modify root growth and architecture in 9311, a traditional *indica* rice cultivar (Figure 3C). β-cyclocitral-treated root systems grew twice as deep as control plants and were significantly narrower (Figure S10). This change in root system architecture depth, was due, in part, to increased primary root growth (Figure S10), but crown roots also grew about 80% deeper when exposed to β-cyclocitral (Figure 3E). This was due both to an increase in crown root length (Figure S10) and a steeper angle of growth (Figure S10). The total number of crown roots per plant also increased by approximately 50% (Figure 3F). The combination of these factors generated deeper, more compact root systems. The overall enhanced root growth caused by β-cyclocitral did not have an obvious effect on shoot mass, which indicates that β-cyclocitral does not have deleterious effects on shoot growth (Figure S11). The added complexity of monocot root systems leads to additional emergent phenotypes that would not have been predicted based on the eudicot studies and reveals new potential roles of β-cyclocitral in root development.

Abiotic stresses such as salinity have a negative effect on plant vigor and root growth. To determine if β-cyclocitral can promote root growth under abiotic stress, we applied it to salt-stressed rice roots (Figure 4A-B). Treatment of seedlings with 50 mM sodium chloride (NaCl) in media significantly decreased root depth compared to control treatment. Root depth could be completely recovered upon co-treatment with β-cyclocitral (Figure 4A-B). In fact, β-cyclocitral had a significantly larger effect on salt-treated plants as compared to unstressed plants (p value ≤ 0.0001, two-way ANOVA). Salt stress also significantly increased the solidity of the root network (Figure S12), measured by calculating the total area of each root divided by the convex area of the root system (33). Increased solidity during salt stress indicates that the plants are producing denser and smaller root systems. Although β-cyclocitral does not affect the solidity of the root system in the absence of salt, it rescues solidity during salt treatment. These results indicate that β-cyclocitral not only enhances root growth during normal conditions but could also shield roots from the harmful effects of salt stress. To test whether this could be an effective treatment in a more natural environment, we performed a salt-stress experiment in soil. Rice grown in a heterogeneous salt-contaminated soil environment had significantly longer roots, on average, when treated with β-cyclocitral (Figure 4C-E). In addition, this treatment increased shoot height, suggesting that it enhanced overall growth. These results indicate that β-cyclocitral treatment could be a beneficial strategy to enhance root growth and plant vigor in agriculture.

**Figure 4.**
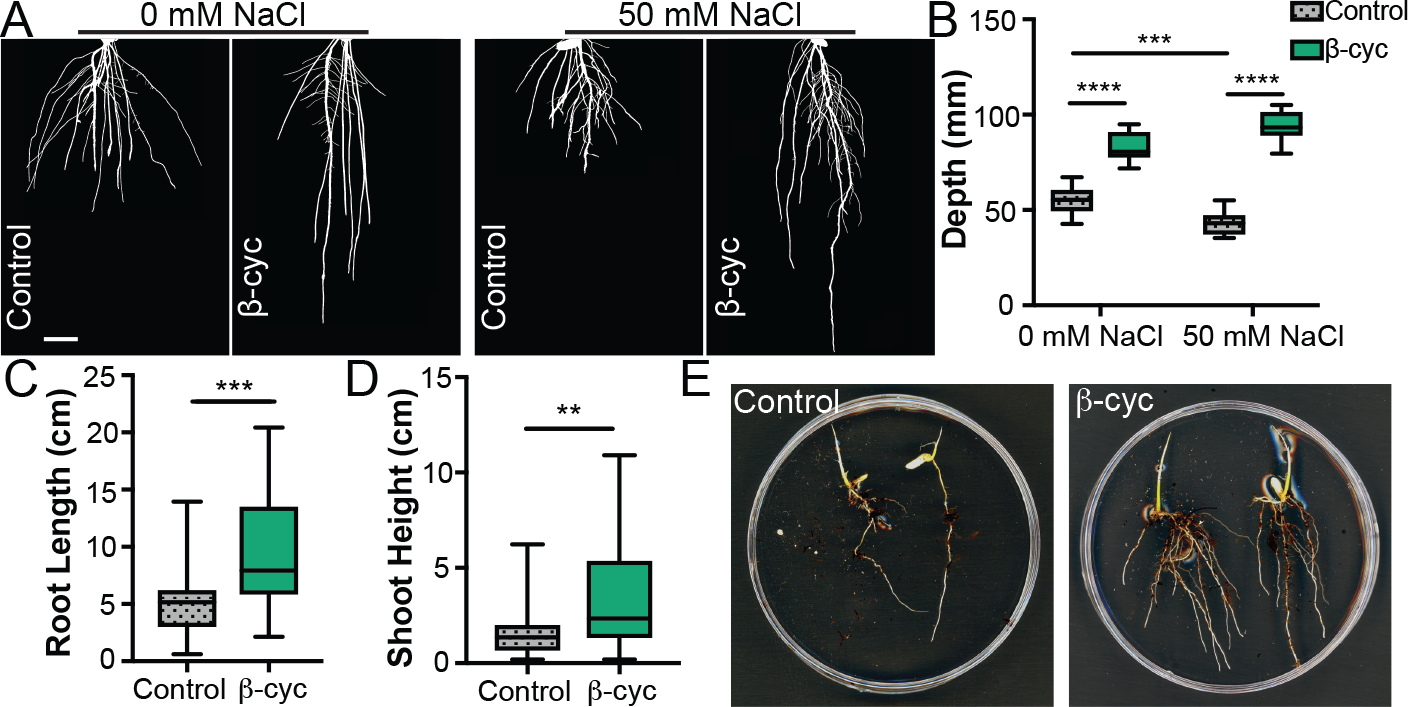
β-cyclocitral promotes rice root growth under salt stress. A) Rice roots treated with β-cyclocitral and grown in gel with 50 mM NaCl (scale bar = 10 mm). B) Root system depth in seedlings treated β-cyclocitral and grown in gel with 50 mM NaCl. C) Primary root length in β-cyclocitral-treated rice plants grown in soil with salt stress. D) Shoot height in β-cyclocitral-treated rice plants grown in soil with salt stress. E) Representative images of rice seedlings grown in salt-contaminated soil with and without β-cyclocitral treatment. The symbols **, ***, and **** indicate p values ≤ 0.01, 0.001, and 0.0001, respectively.

## Conclusion

Through a sensitized chemical genetic screen, we identified β-cyclocitral as a naturally occurring β-carotene derived apocarotenoid, which regulates root architecture in monocots and eudicots. β-cyclocitral is a natural compound that is inexpensive, active at low concentrations, and can be applied exogenously, making it a promising candidate for agricultural applications. Using *Arabidopsis* as a model system, we found that β-cyclocitral increases primary and LR growth by inducing cell divisions in root meristems. β-cyclocitral additionally has conserved effects as a root growth promoter in tomato and rice. In rice, β-cyclocitral also affects other aspects of root architecture, including the numbers and gravity set-point angle of roots. In salt-stressed rice roots, β-cyclocitral significantly promotes root and shoot growth. These results indicate that β-cyclocitral is a natural compound that could be a valuable tool to improve crop vigor, especially in harsh environmental conditions.

## Acknowledgments

The authors wish to thank Jingyuan Zhang for technical support provided for the duration of this project; Michel Havaux and Stefano D’Alessandro for helpful discussions; Dolf Weijers, Cara Winter, Colleen Drapek, and Isaiah Taylor for critical reading; and Linxing Yao at the Proteomics and Metabolomics Facility at Colorado State University for experimental assistance.

## Author contributions

AJD and PNB conceived of this study. AJD, KL, JM, KJ, MM and **PNB** designed methodology, performed experimental investigations, and analyzed data. AJD and **PNB** wrote the original drafts of this paper, with review provided by KL, MM, JD, JM, KJ, and SA. JD, JM, KJ, and SA provided experimental and conceptual support during the revisions.

## Data and materials availability

All data is available in the main text or the supplementary materials.

## Supplementary Materials

Materials and Methods

Figures S1-S8

Table S1

References (29-32)

